# Solid/liquid coexistence during aging of FUS condensates

**DOI:** 10.1101/2022.08.15.503964

**Authors:** Yi Shen, Anqi Chen, Wenyun Wang, Yinan Shen, Francesco Simone Ruggeri, Stefano Aime, Zizhao Wang, Seema Qamar, Jorge R. Espinosa, Adiran Garaizar, Peter St George-Hyslop, Rosana Collepardo-Guevara, David A. Weitz, Daniele Vigolo, Tuomas P. J. Knowles

**Affiliations:** Yusuf Hamied Department of Chemistry, University of Cambridge, CB2 1EW Cambridge, UK; School of Chemical and Biomolecular Engineering, The University of Sydney, NSW 2006, Australia; The University of Sydney Nano Institute, The University of Sydney, NSW 2006, Australia; John A. Paulson School of Engineering and Applied Sciences, Harvard University, Cambridge, MA 02138, USA; Department of Physics, Harvard University, Cambridge, MA 02138, USA; Laboratory of Organic Chemistry, Wageningen University, 6708 WE, the Netherlands; Physical Chemistry and Soft Matter, Wageningen University, 6708 WE, the Netherlands; Molecular, Macromolecular Chemistry, and Materials, ESPCI Paris, CNRS, PSL University, 75005 Paris, France; Cambridge Institute for Medical Research, Department of Clinical Neurosciences, University of Cambridge, CB2 0XY Cambridge, UK; Cavendish Laboratory, University of Cambridge, CB3 0HE Cambridge, UK; Department of Medicine, Division of Neurology, University of Toronto and University Health Network, ON M5T 0S8 Toronto, Canada; Department of Genetics, University of Cambridge, Cambridge, CB2 3EH Cambridge, UK; Wyss Institute for Biologically Inspired Engineering, Harvard University, Boston, MA 02115, USA; School of Biomedical Engineering, The University of Sydney, NSW 2006, Australia

## Abstract

A wide range of macromolecules undergo phase separation, forming biomolecular condensates in living cells. These membraneless organelles are typically highly dynamic, formed in a reversible manner, and carry out important functions in biological systems. Crucially, however, a further liquid-to-solid transition of the condensates can lead to irreversible pathological aggregation and cellular dysfunction associated with the onset and development of neurodegenerative diseases. Despite the importance of this liquid-to-solid transition of proteins, the mechanism by which it is initiated in normally functional condensates is unknown. Here we show, by measuring the changes in structure, dynamics and mechanics in time and space, that FUS condensates do not uniformly convert to a solid gel, but rather that liquid and gel phases co-exist simultaneously within the same condensate, resulting in highly inhomogeneous structures. We introduce two new optical techniques, dynamic spatial mapping and reflective confocal dynamic speckle microscopy, and use these to further show that the liquid-to-solid transition is initiated at the interface between the dense phase within condensates and the dilute phase. These results reveal the importance of the spatiotemporal dimension of the liquid-to-solid transition and highlight the interface of biomolecular condensates as a key element in driving pathological protein aggregation.

---

Liquid-liquid phase separation (LLPS) is a result of spontaneous decomposition of a uniform solution into two aqueous phases: a dense phase rich in molecules and a dilute one depleted in molecules. While LLPS takes place for many polymeric systems, its central role for functional biology was first recognized in germline P granules (*1*). Subsequently, many other membraneless organelles have been identified, contributing to essential biological processes, such as DNA damage response (*2*), nuclear trafficking (*3*) and signal transduction (*4*). The dynamic nature of the condensates is the key characteristic required for favoring chemical reactions within the granules and molecular exchange with the dilute phase outside. However, the protein-rich phase of LLPS has the potential to also enhance aberrant protein-protein interactions, and therefore increases the propensity of aggregation that can lead to the formation of irreversible pathological structures(*5, 6*). Such irreversible aggregation initiated from initially reversible biomolecular condensates plays an important role in neurodegeneration; for instance, α-synuclein aggregates, linked to Parkinson’s disease pathogenesis, have been found to nucleate through from a condensate precursor *in vivo* (*7*). Furthermore, tau, a protein associated with Alzheimer’s disease, forms irreversible intracellular deposits in disease which originate from initially reversible condensate states (*8, 9*). Similarly, Transactive response DNA-binding protein of 43 kDa (TDP-43) and the fused in sarcoma (FUS) proteins, connected to amyotrophic lateral sclerosis and frontotemporal lobar degeneration (ALS/FTLD), also form fibrillar aggregates stemming from reversible and functional condensates (*5, 10–12*). It is therefore of key importance to understand the determinants of the liquid-to-solid transition of condensates, as they have direct implications for the emergence of pathological states through incorrectly assembled proteins. To date, the spatiotemporal dimension of this liquid-to-solid transition has not been explored in detail and condensates are generally considered as homogenous droplets rich in molecules (*1, 11, 13, 14*) which undergo gelation. Here, however, by deploying advanced characterization approaches, we show that condensates do not uniformly convert to a solid gel, but rather that liquid and gel phases co-exist simultaneously within the same condensate, resulting in highly inhomogeneous structures. Our results also suggest that there is a propensity of the liquid-to-solid transition to initiate from the interface and propagate towards the center of the condensates. These discoveries of the intricate role of spatiotemporal dynamics highlight the complexity in the patterns associated with the liquid-to-solid transition and reveal the interface of the condensates as a particularly crucial element in this process.

We focus in our study on a key protein implicated in neurodegenerative diseases, FUS (Fused in Sarcoma). For imaging purpose and consistency within the experiments, we use GFP-FUS throughout the study. By using confocal imaging, we observe the phase transitions of condensates, from liquid droplets to liquid-solid coexisting gel to fibrillar solid (Fig. 1A). By focusing on the density from the distribution of the fluorescence intensity, we can detect changes at early time in the spatial distribution of molecules as a function of aging of the condensate (Fig. 1B). During the first 30 minutes, the protein molecules are uniformly distributed in the condensates. Signs of spatial inhomogeneity can, however, be detected after few hours. After 24 hours of incubation, the condensates become less homogeneous and present a weak core-shell structure (Fig. 1BC). Moreover, through confocal 3D scanning, the condensate shows a loosely packed network (Mov. S1). After further aging for 5 days, clearly defined filamentous structures form around the condensates (Fig. 1D), characteristics of the irreversible aggregation, representing the solid end state of the transition, consistent with the previous studies (*5*).

**Fig. 1.**
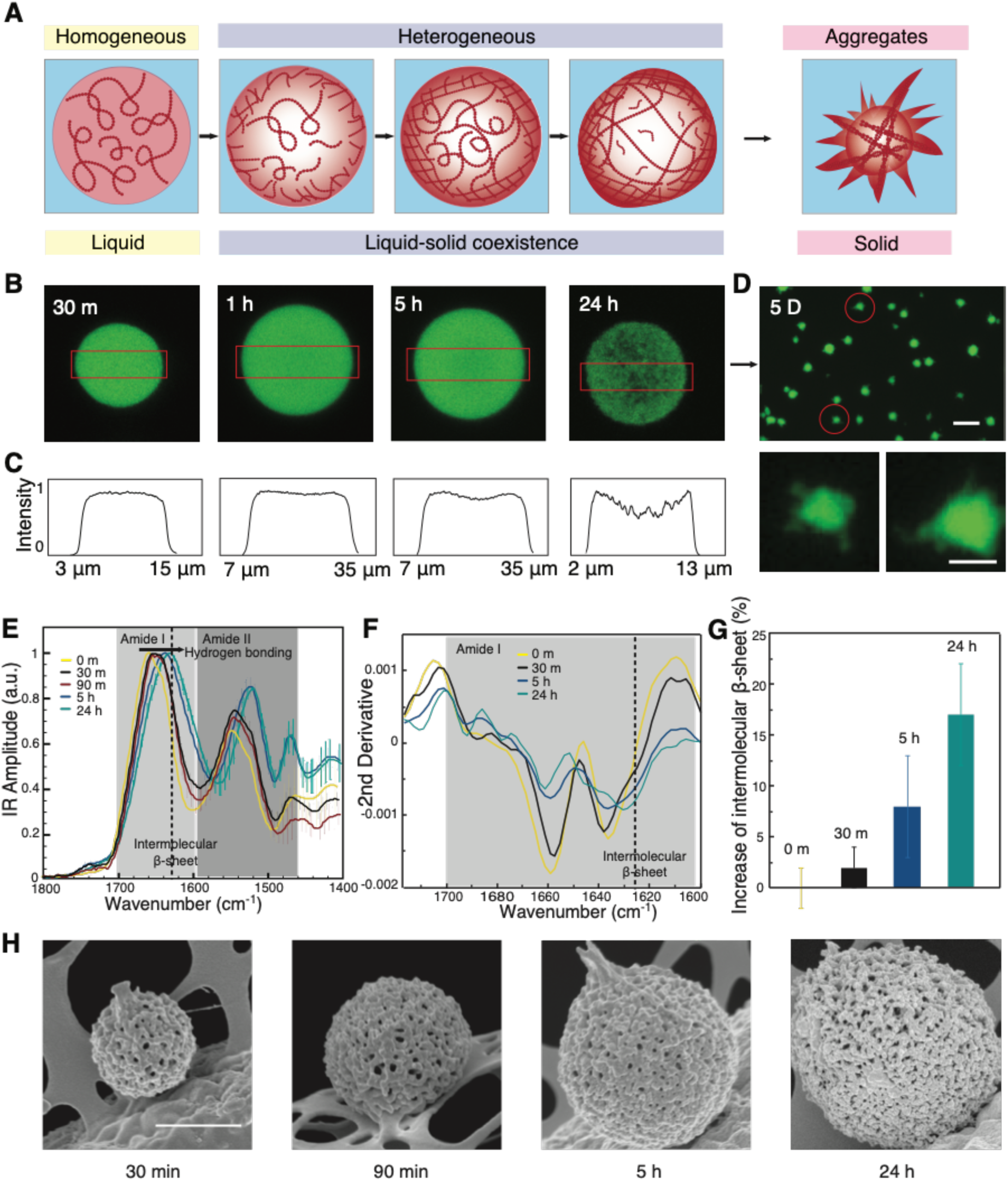
Spatially inhomogeneous liquid-to-solid transition. A. The FUS protein undergoes liquid-liquid phase separation in response to changes in ionic strength. During subsequent maturation, the distribution of the molecules within the condensates changes from homogenous to heterogeneous B. Confocal images of the density distribution within a condensate during maturation at 30 min, 1h, 5h and 24h. C. The fluorescence intensity profile from confocal microscopy at the center plane of the condensates (the evaluated area is indicated by red boxes in B). D. Fluorescence images of fibrillar aggregates formed at the final stages of the liquid-to-solid transition after maturation for 5 days (Scale bars are 20 and 5 μm in top and bottom panels). E. Bulk IR absorption spectra and F. their second derivatives of condensates with different aging time. G. Relative increase of intermolecular β-sheet with aging time, error bar: confidence interval deriving from the integration algorithm of the second derivative. H. SEM vimages of condensate with different maturation time, scale bar 1 μm.

To characterize the molecular level changes associated with the liquid-to-solid transition, we further perform bulk infrared spectroscopy (FTIR) on the liquid-liquid phase separated samples. The spectra show a net shift of both amide band I and amide band II towards lower wavenumbers as a function of the incubation time from 30 minutes to 24 hours, thus demonstrating the formation of hydrogen bonding associated to intermolecular β-sheet (*15–17*) (Fig. 1E, Fig. S1). To further evaluate the secondary and quaternary structural changes that the LLPS droplets undergo as a function of the incubation time, we perform a second derivative analysis of the amide band I (*18, 19*). The second derivative spectra show a continuous decrease of α-helix and random coil structure (1660-1640 cm^-1^) versus an increase of intermolecular β-sheet structure (1630-1620 cm^-1^) as a function of the incubation time (Fig. 1F). The relative increase of the intermolecular -sheets with aging time compared to the native droplets is evaluated by direct integration of the area of the structural contributions in the second derivative spectra and is summarized in Figure 1G. The increase of intermolecular β-sheet and the continuous shift at lower wavenumbers demonstrate the formation of longer intermolecular β-sheets and a denser network of hydrogen bonds, underlying the mechanism of the LST (*15, 16*). We next observe these condensates under SEM after flash freezing and drying. We find well-defined fibrillar networks covered all over the condensates (Fig. 1H). Thus, these results show a liquid-to-solid transition of the condensates, contributed by the intermolecular β-sheets formation with a fibrillar network.

After having characterized the overall liquid-to-solid transition, we focus on probing the spatiotemporal dynamics. First, we probe for heterogeneity developing within the condensates during the liquid to solid propagation by conducting a set of dissolution experiments on condensates (Fig. 2). To test the formation of irreversible interactions between protein molecules in the condensate, we add high concentration KCl solution to the condensates at different points during the aging process. Relatively fresh condensates (aged for 1h) rapidly dissolve upon the addition of 1M KCl solution, as predicted by phase diagrams measured in previous studies (*20*). By contrast, when 1M KCl solution is added to condensates aged for 72h, no overall dissolution is observed, indicating the formation of irreversible interactions. Only a minor drop of fluorescence intensity is seen in these samples, showing the release of a residual amount of un-gelled monomers. To investigate the intermediate gelation stages, we repeat the dissolution experiment with the 24h and the 48h samples. Surprisingly, when 1M KCl is added to the 24h condensates, the center of the condensate dissolves, leaving behind a thin shell of gelled material. When the 48h condensates are subjected to high concentration KCl, the center dissolves as seen in the 24h sample. However, a thicker shell is left behind (Fig. 2AB). From confocal 3D reconstructed images, a clear shell structure is observed (Fig. 2C). Taken together, these results suggest the gelation of the condensates initially happens at the interface and propagates inward with time.

**Fig. 2.**
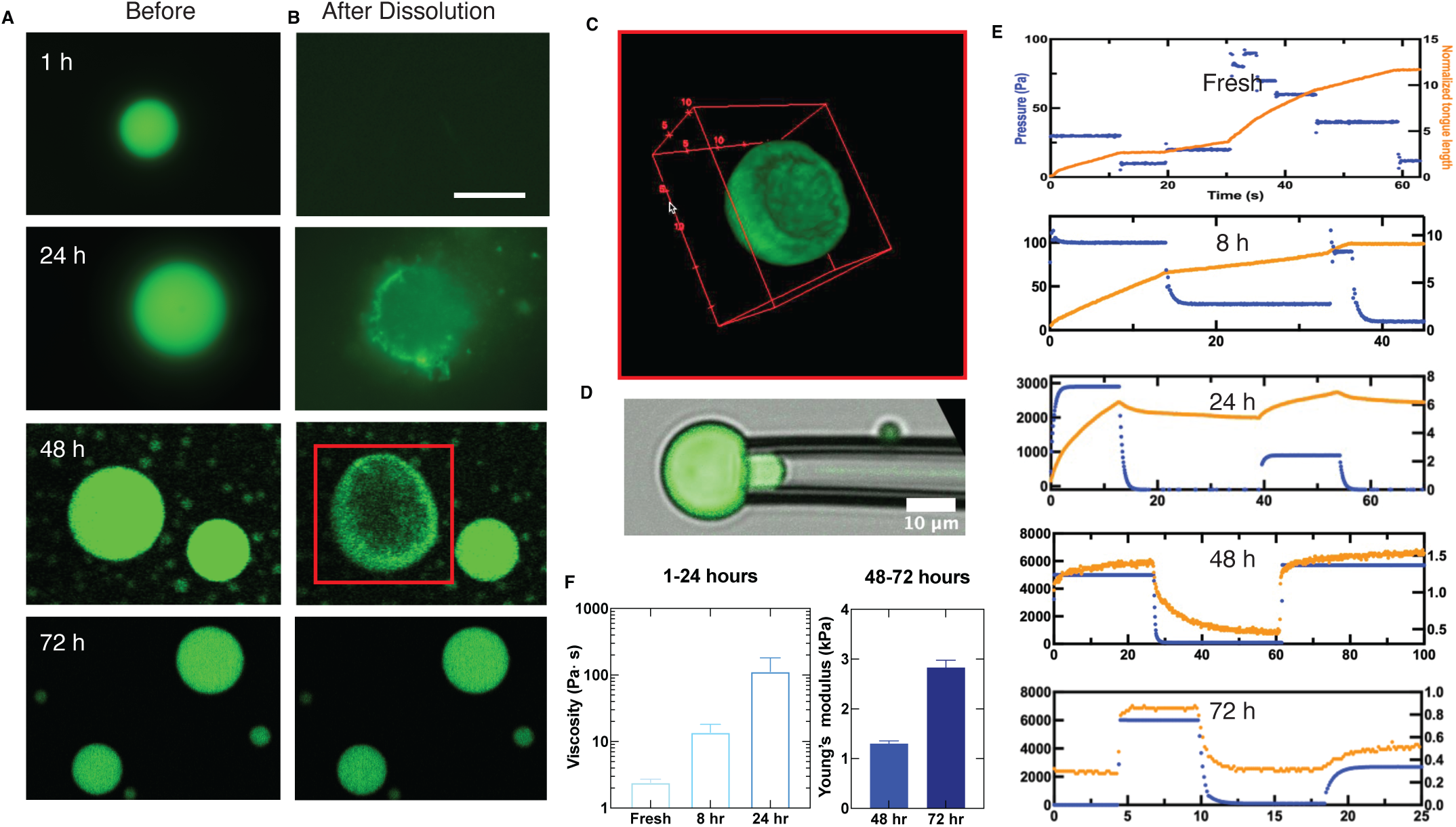
Dissolution and micromechanics of condensates. A. Condensates incubated at 25°C for 1 h, 24 h and 45 C for 24 h. B. Dissolution of condensates monitored after introduction of a solution 1M KCl in volume ratio 1:10 for 5 minutes. C. 3D reconstructed image of a condensate incubated for 48h after KCl partial dissolution from KCl treatment as in B. D. Micropipette aspiration of a condensate E. the relationship between tongue length and the elapsed time under different pressure during the maturation of the condensates from fresh to 72 h. F. The increase of viscosity of condensates from fresh to 24h (left) and the increase of elasticity of condensates incubated for 48h and 72h evaluated form the data in E.

To connect these changing spatial patterns to variations in the mechanical properties of the condensates during aging we use micropipette aspiration experiments (*21*). The condensates are aspirated using a glass micropipette with an inner diameter of 4 – 10 μm. Several negative pressure steps are applied using a vacuum controller, and the length of the condensate drawn into the glass pipette is measured. This approach determines both the strain and the stress associated with the deformation and therefore probes the rheological response of individual condensates. Under constant-pressure aspiration, a liquid-like condensate flows into the glass pipette with a constant velocity, resulting in a linear increase of the length of the tongue as a function of elapsed time. The resultant strain rate is proportional to the applied pressure and viscosity of the liquid-like condensate. By contrast, a solid-like condensate deforms rapidly when the pressure changes but then reaches a constant value. In this case, the strain, determined by the tongue length, is proportional to the stress, determined by the applied pressure, providing a measure of the elastic modulus of the condensate. By using this method, the condensates are probed at different aging stages. Upon formation, the fresh FUS condensate is a liquid with a viscosity of order 1 Pa×s. In the following 8 h, the condensates remain liquid-like with gradually increasing viscosity. After 24 h, we measure an increase in the viscosity of ca. 50 times its initial value, although the elasticity is still too small to measure precisely. After 48h, the condensates exhibit an elasticity-dominant mechanical response with a viscoelastic creep behaviour. The elastic modulus is ∼ 1 kPa. After 72h, the elastic modulus increases to ∼ 3 kPa. We attribute the dramatic change of the mechanical properties of FUS condensates to the formation of a solid gel network within them.

From confocal 3D scanning, it is observed that there is a heterogeneous organization inside the condensates after 24 hour incubation, where a weak gel network structure has developed (Fig. 3A). However, it is less clear for the condensates aged for less time (Fig. 3B). To unveil and quantify internal heterogeneities of the condensates we develop and apply a Spatial Dynamic Mapping (SDM) approach. The condensates exhibit refractive index fluctuations through thermal motion. Thus, light travelling through a condensate forms patterns that vary over time which can be analysed to extract information regarding their internal dynamics and hence, potentially, their internal structure. We divide the condensates into small regions of interest (ROI) and we evaluate the spatial variance in real space, *σ*^2^(Δ*t*), from each ROI within each frame. We use a fast camera to record bright field images and can thus acquire long sequences of images at high frame rate. This gives us the ability to explore simultaneously fast (acquiring at high frame rates) and slow dynamics (by acquiring for long time). We are thus able to extrapolate the characteristic decay time from the exponential behavior of *σ*^2^(Δ*t*) and we use this to create the spatial map in Fig. 3C. From these maps it is possible to discover the coexistence of areas within the condensate that exhibit a liquid-like (characterized by a short decay time) and solid-like (longer decay time) behavior (detailed calculation and calibration can be found in SI). In this study, SDM can reliably discriminate between characteristic decays spanning 3 orders of magnitude, ranging between one and a thousand seconds, within the same condensate. We further perform SDM in the middle plane and at the bottom plane of a condensate exploiting the ability of SDM to extract 3D information and find that the bottom plane shows slower dynamics with higher heterogeneity. We plot the radial distribution of the characteristic time as a function of the distance from the center for both the middle and bottom planes. The middle plane demonstrates slower dynamics close to the interface of the condensate (Fig. 3D). These results confirm the core-shell structure of the condensates.

**Fig. 3.**
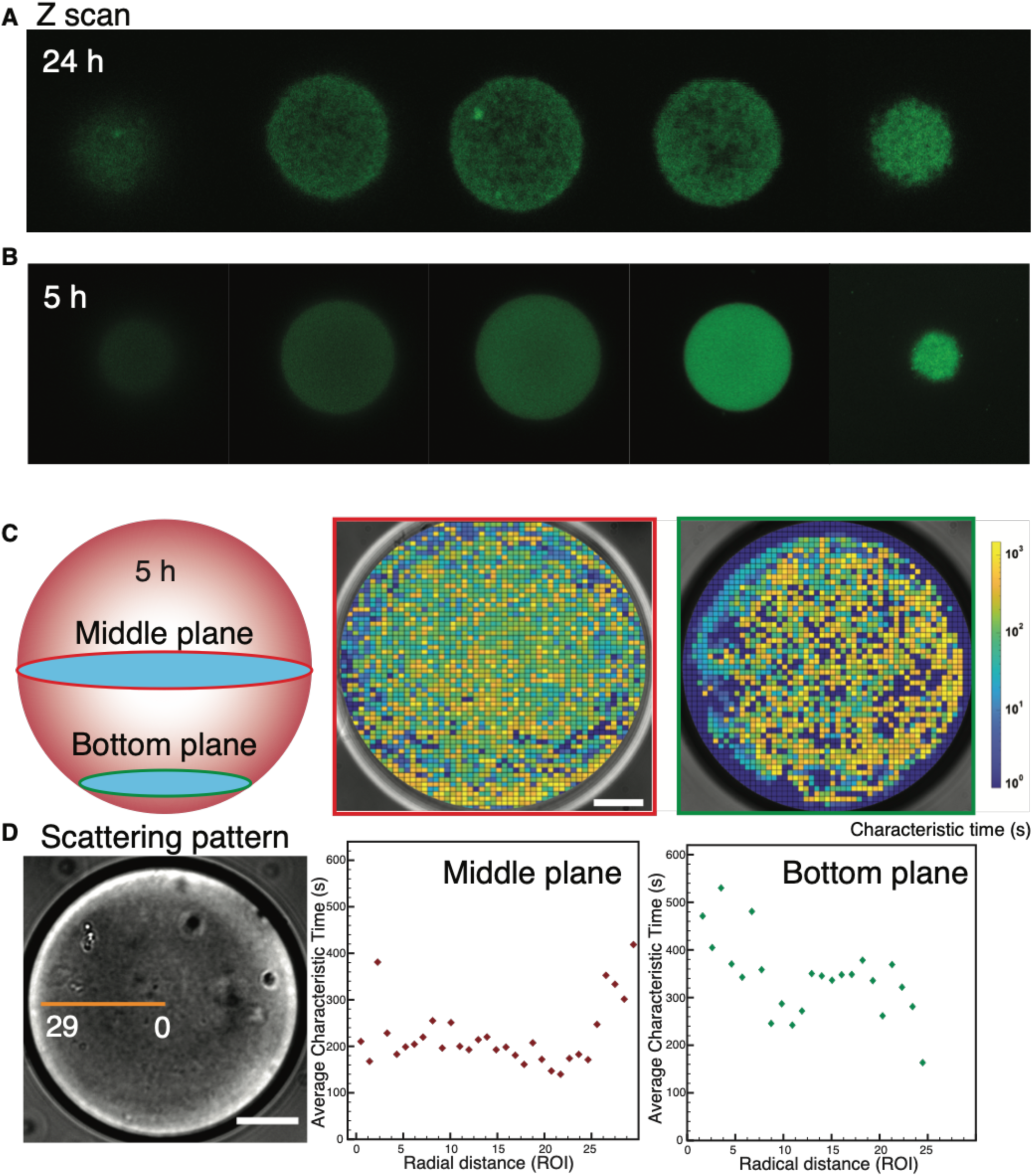
Confocal z-scan of condensates incubated for A. 24h and B. 5h. C. Dynamic Spatial mapping of the characteristic correlation time of the condensate at middle and bottom planes with aging time 5 h, scale bar 20 μ*m*. D. Radial distribution of the characteristic time from the center of the condensate to the edge within a condensate aged for 5h.

To achieve a more sensitive measure of the fluctuations of the structure in the condensate, we develop a new optical technique, reflective confocal dynamic speckle microscopy. We use a laser scanning confocal microscope to image the backscattered light from a plane near the center of the condensate. Surprisingly, when the confocal pinhole is adjusted to match the Airy disk of the image, a speckle pattern is observed, as shown for a condensate after 24 h in Fig. 4A. This pattern fluctuates in time as successive images are collected at a rate of 20 Hz. We calculate the temporal correlation of each pixel and average over an area corresponding to the speckle size, which is determined from a spatial correlation of the image. These temporal correlation functions exhibit a decay to a constant value. Since we are imaging a speckle pattern, the predominant origin of the fluctuations is relative motion of scatterers, within the diffraction-limited volume detected with the confocal, in a direction parallel to the scattering vector. We emphasize that, although we are performing a scattering experiment, we are nevertheless imaging the sample with a confocal microscope; therefore, the information provided by this experiment is sensitive to very small motions, which cause the speckle intensity to vary, but is still localized to the diffraction volume. We can, therefore, determine the local dynamics with high spatial resolution within the condensate. The correlation functions are well described as a stretched exponential decay to a constant plateau value, 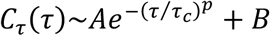, where *A* and *B* are constants, *p* is the stretching exponent and *τ*_*c*_ is the decay time, as shown by the logarithmic plots of *C*_*τ*_ (*τ*) as a function of *τ*^*p*^ in Fig. 4b. We use *p* = 0.4, although the best fit varies slightly, between 0.3 ≤ *p* ≤ 0.5, for all the speckles in the image. Similarly, the plateau value varies across the image, with many speckles decaying completely (*B* = 0) while others having plateaus as high as *B* = 0.8. This behavior is consistent with scattering from a solid, gel-like network: the stretched exponential decay reflects relaxation of the imaged volume due to fluctuations of all length scales that contribute to the relaxation (*22, 23*). The plateau reflects solid-like behavior.

**Figure 4.**
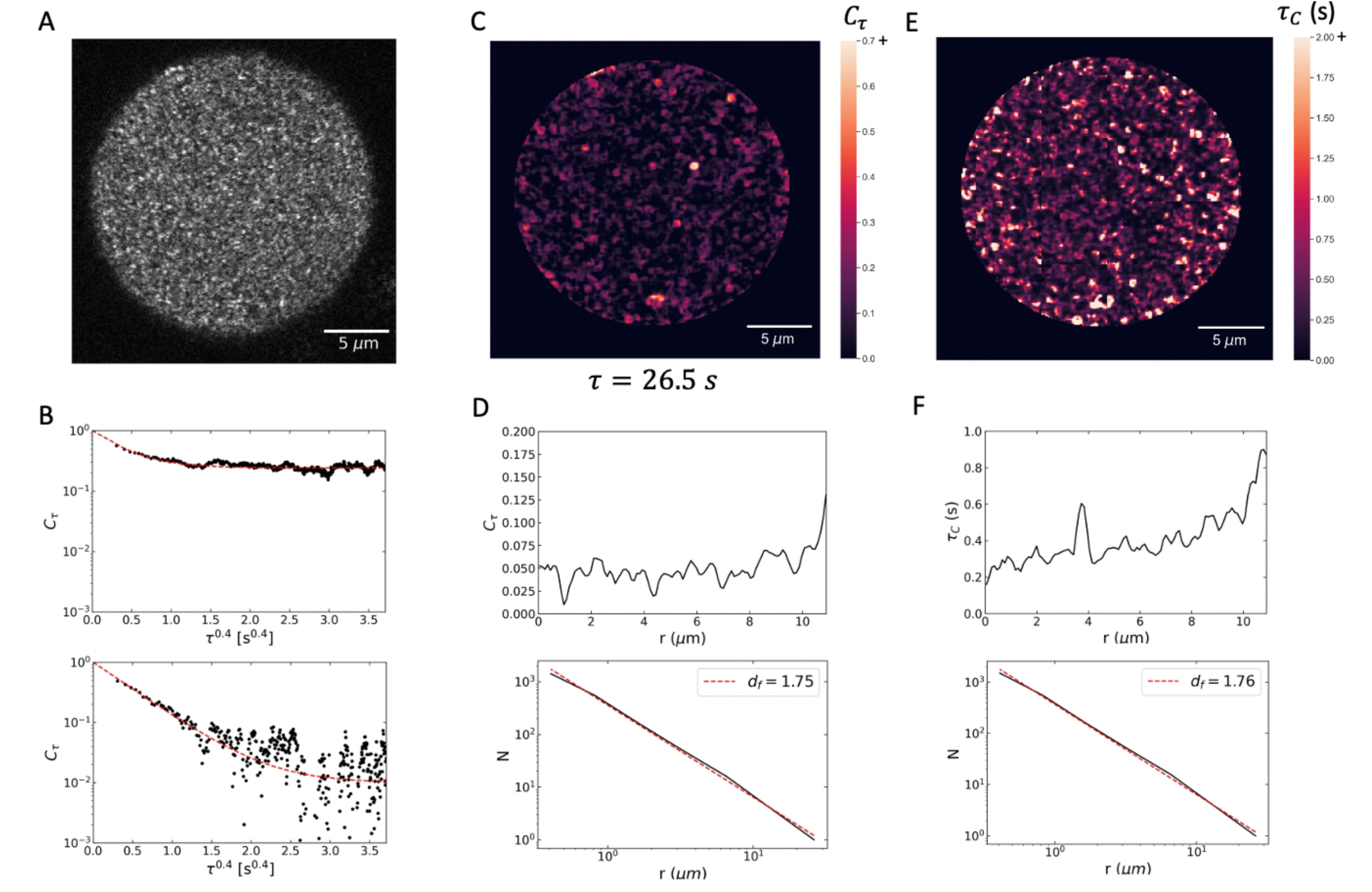
Confocal backscattering speckle fluctuation analysis: A. Image of speckles from a 24h condensate. B. Temporal correlation function of two speckles showing stretched exponential decay to a plateau. The upper correlation function decays to a plateau, indicating a solid, while the plateau of the lower correlation function is at the noise level of the measurement. C. Spatial map of the plateau region of the temporal correlation functions D. Radial distribution of the value of *C*_*τ*_(*τ*) at *τ*= 26.5 s, well into the plateau region (top) and box counting analysis of the fractal dimension of the plateau regions using a threshold of *C*_*τ*_ = 0.10. The structure is fractal with *d*_*f*_ *≈* 1.75. E. Spatial map of decay times of the stretched-exponential correlation functions. F. Radial distribution of decay times exhibiting pronounced increase at outer edge (top) and box counting analysis of decay times greater than 0.5 s (bottom). The structure is fractal with *d*_*f*_ *≈* 1.76. The corresponding fractal dimension for the three-dimensional condensate is *d*_*f*_ ≈ 2.75.

The spatial dependence of these correlation functions provides important new insight into the structure and dynamics of the condensate. The plateau in the correlation functions is a direct measure of solid-like behavior, and crucially it does not exhibit a uniform value throughout the condensate. Instead, there are regions that are solid-like and regions that are liquid-like, suggesting that there is a further spatial phase separation of the solid portion with respect to the remaining liquid phase. To investigate the structure of this solid-like region, we plot the value of *C*_*τ*_ (*τ*) at a lag time of *τ*= 26.5 s in Fig. 4C. A radial average of the plateau values shows a distinct increase at the outer edge of the condensate, supporting the view that the liquid-solid transition begins at the outer edge, as shown in the upper plot in Fig. 4D. Perhaps even more interesting is that the structure within the condensate is not homogeneous, but instead is quite heterogeneous. To quantify the spatial distribution of solid-like regions, we set a threshold of *C*_*τ*_ (*τ*) = 0.10 and use box-counting to determine the spatial structure. It is well described as a fractal as shown by the power-law dependence plotted in Fig. 4F. The fractal dimension of the image is *d*_*f*_ ≈ 1.76 as shown by the dashed line in Fig. 4F; however, since this is a two-dimensional slice through a three-dimensional image, the fractal dimension of the condensate is *d*_*f*_ ≈ 2.26. Similar heterogenous structure is found in the spatial distribution of the decay times as shown in Fig. 4E. The radial average of *τ*_c_ exhibits a pronounced peak at the outer edge of the condensate, as shown in the upper plot in Fig. 4F. Setting a threshold at *τ*_c_ = 0.5 s and performing a box-counting analysis again shows a fractal structure with essentially the same value of *d*_*f*_ as shown in the lower plot in Fig 4F. These results are strong evidence of a heterogeneous structure within the condensate, reflecting a solid, gel-like network in a more liquid-like background. We can estimate the elastic modulus of this network from the plateau value of the correlation functions. The dimension of each fluctuation element contributing to the speckle is ~*q*^−1^, where *q* is the scattering vector. We take the value of the plateau as 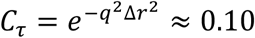. This gives an elastic modulus of *G*′ ∼ *q*^3^*k*_*B*_*T*, where *k*_*B*_*T* is the thermal energy. From this, we obtain *G*’~120 Pa. Thus, the analysis of the backscattered speckles supports the view that the condensate highly heterogenous, reflecting a second spatial phase transition, leading to the formation of a very weak solid gel-like network.

To explore the connection between molecular level interactions and the observed liquid/solid co-existence during the liquid-to-solid transition of condensates, we conduct molecular simulations with a dynamical algorithm coupled to a coarse-grained model for FUS proteins in explicit solvent, which we developed recently using a multiscale approach (*24*). Our dynamical algorithm describes the nonequilibrium process of condensate aging by introducing dissipation via non-conservative inter-protein interactions. Specifically, we first perform Direct Coexistence simulations, starting from a well-mixed solution of FUS molecules at subcritical conditions (Fig. 5B), to obtain an equilibrium condensate in coexistence with a diluted liquid. We then turn on the dynamical algorithm. As time progresses, our dynamical algorithm triggers disorder-to-order transitions (i.e., from disordered to inter-protein β-sheets) within the prion-like domain (PLD) of individual FUS molecules (Figure 5a). Protein structural transitions are recapitulated by modulating the interaction strength of PLD–PLD interactions—according to atomistic potential of mean force simulations (*24*)—and implemented only when high local fluctuations of protein densities emerge within the condensate. As a result, during our aging simulations, FUS condensates exhibit gradual accumulation of strongly binding and locally rigid inter-protein β-sheets (Fig. 5C). As seen in Fig. 5C, the number of structural transitions rapidly increases over time, and then plateaus as the lower density core of the condensate consolidates (Fig. 5D), showing the transitions are disfavored at lower protein concentrations but favored at the condensate-solvent interphase (Fig. 5E). The simulations recapitulate the co-existence of liquid and solid phases within a single condensate during the gelation process. Moreover, consistent with our experiments, at the molecular scale of the interphase of the condensate we observe that inter-protein β-sheets form preferentially at the interphase because it is there where larger fluctuations in local protein density occur (Fig. 5E) – such density fluctuations are precisely the ones that seed the structural transitions. Finally, Fig. 5F shows the mean-square displacement of proteins with structured PLD is much smaller compared to proteins with unstructured protein in a condensate. Combining with Fig. 5E, the results indicate that the outer shell of the condensate, where the proteins have undergone the transitions, is more akin of a gel than a liquid (yellow curve), and the inner core remains liquid (red curve). The results also provide support for the development of the internal structure of the condensate, which exhibits a localized liquid-solid transition similar to that in Fig. 5E.

**Figure 5.**
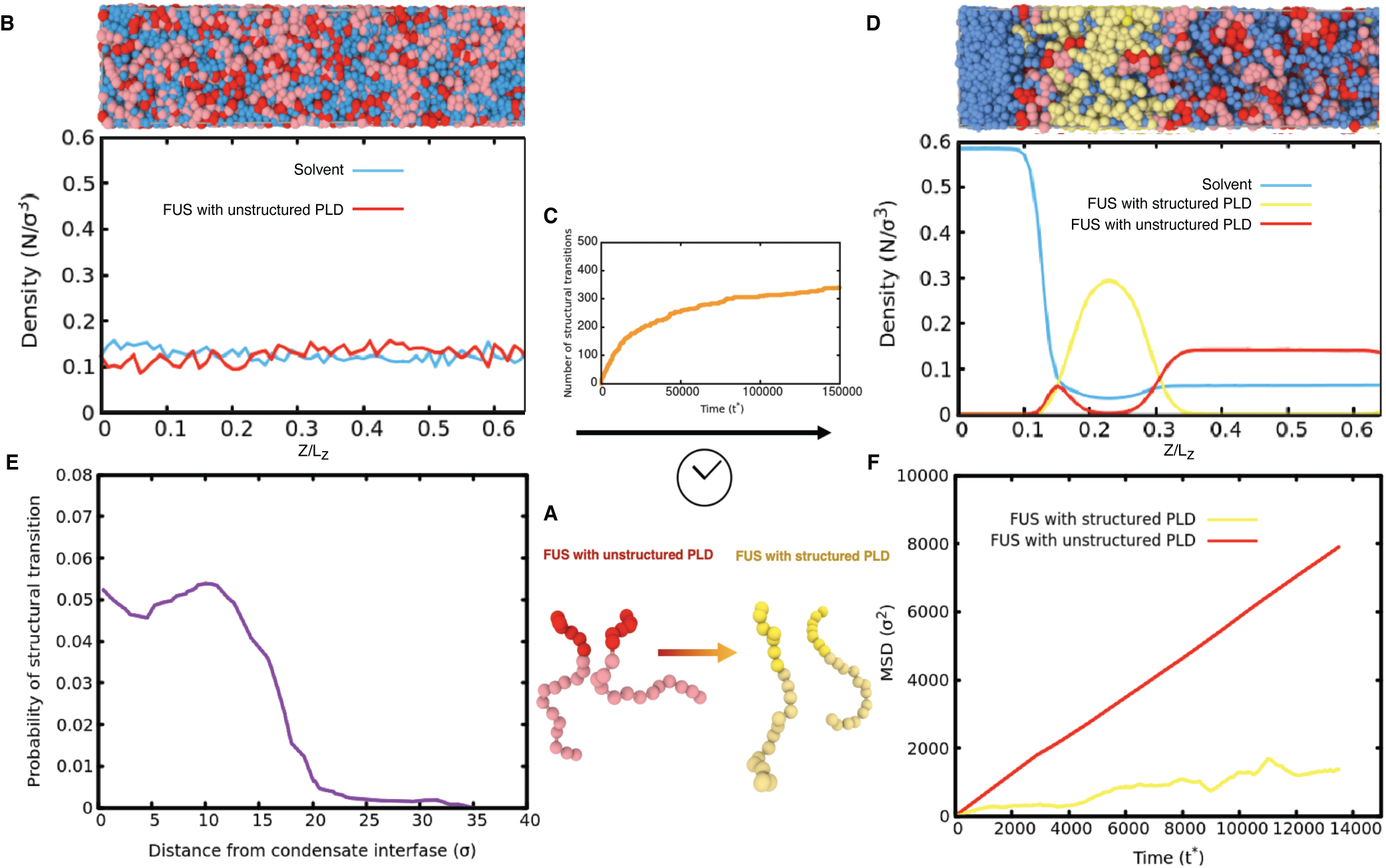
Molecular simulations with a dynamical algorithm describe the emergence of spatially inhomogeneous gel phase co-existing with a liquid phase during aging. A. Minimal coarse-grained models for FUS with fully disordered PLDs (Red) and with structured PLDs (i.e., with inter-protein β-sheet elements in the PLDs) (yellow). B. Top: Snapshot of initial state in our direct coexistence simulations depicted a well-mixed solution of FUS proteins with fully unstructured PLDs (Red) and explicit solvent (Blue) Bottom: Density profile (in reduced units) showing an initial homogeneous distribution of FUS species and explicit solvent across the long side of the simulation box. C. Number of structural transitions in FUS-PLD domains as a function of simulation time (t_*_). D. Top: Snapshot of direct coexistence simulations using the dynamical algorithm at a steady state after structural transitions have saturated. Bottom: Density profile (in reduced units) of FUS species and explicit solvent across the long side of the simulation box estimated over the coarse-grained steady state. E. Probability of emergence of a structural transition inside the FUS condensates during aging as a function of the distance from the interphase. F. Mean squared displacement of FUS proteins with structured PLDs versus FUS proteins with fully disordered PLDs.

In summary, liquid-liquid phase separation is increasingly well understood as an essential process with crucial biological functions. By contrast, the mechanisms of liquid-to-solid transition of condensates, underlying the formation of pathological protein aggregates, have remained more challenging to elucidate. In this study, we develop and deploy a set of advanced imaging techniques to probe the onset and development of a solid phase within liquid FUS condensates at multiple time and length scales. We discover that there are multiple dynamical time scales within a single condensate at all levels of maturation, indicating the co-existence of liquid and gel network within a condensate. We further unveil that the material properties of the condensates transition from liquid-like to solid-like from the interface to the core as a function of the maturation time. A core-shell structure starts to form after 24 hours of incubation time and the growth of the solid shell propagate towards the center until the whole condensate becomes gel. Our findings unveil the spatiotemporal dimension of the liquid-to-solid transition of condensates and may suggest new interventions to mitigate this transition when it occurs in driving such systems from physiological to pathological.

## Data availability

All the relevant data are included in the manuscript and supplementary information. More detailed protocols, calculation and analysis are available from the authors upon request.

## Acknowledgements

This work is supported by the Newman Foundation, the Wellcome Trust, ERC, Alzheimer Association Zenith, ALS Canada–Brain Canada, Canadian Institutes of Health Research and the Cambridge Centre for Misfolding Diseases. We thank Cambridge Advanced Imaging Centre and K.H. Muller for help with flash-freezing and SEM imaging.

## Author Contribution

Y.S., D.V. and T.P.J.K. conceived and designed the study. Y.S., A.C., W.W, S.A., Y.S., F.S.R., D.V. performed the experiments and analyzed the data. A.G. J.R.E. and R.C-G designed and developed the simulations methods. A.G. and J.R.E performed the simulations and analyzed the simulation data. Z.W. helped with image analysis. S.Q. produced the protein. D.W., R.C-G. and P.S.G.-H advised the study. All authors contributed to the writing of the manuscript.

