## Supplementary material for "Solid/liquid coexistence during aging of FUS condensates": Material methods and supplemental tables and figures

### Materials and Methods

*FUS condensates sample preparation*

FUS protein used in this study was GFP (Green Fluorescent Protein) labelled. The purification method has been described in previous studies(*1–3*). The final purified protein was stored in the buffer with 50 mM Tris, 1 M KCl, 1 mM DTT, 5% glycerol, pH 7.4 at concentration 25 – 35 $\mu$M. The purity of the protein was greater than 95%.

FUS protein stock was mixed with MiliQ water in a volume ratio of 1:19 to induce the LLPS. The final FUS and KCl concentrations were 1.75 – 1.25 $\mu$M and 50 mM, respectively. FUS condensates were then incubated at room temperature for 30 min, 90 min, 5 h, 24 h for aging. For DDM, large condensates were chosen for better resolution.

*Confocal Imaging*

Confocal microscope TCS SP8, Leica Microsystems with a 488-nm laser Argon and ×60 oil-immersion objective were used for imaging. Z-scan was performed at 0.1% power of the 488-nm laser with 100 nm step between the scanning slides.

*Fourier-transform infrared spectroscopy (FTIR)*

Attenuated total reflection infrared spectroscopy (ATR-FTIR) was performed using a Bruker Vertex 70 spectrometer equipped with a diamond ATR element. The resolution was 4 $cm^{-1}$ and all spectra were processed using Origin Pro software. The spectra were averaged (3 spectra with 256 co-averages), smoothed applying a Savitzky-Golay filter ($2^{nd}$ order, 9 points) and then the second derivative was calculated applying a Savitzky-Golay filter ($2^{nd}$ order, 11 points).

Relative secondary and quaternary organization was evaluated by integrating the area of the different secondary structural contribution in the amide band I, as previously described(*4*, *5*).

*Scanning Electron Microscopy (SEM)*

FUS protein condensates were aged and sampled at 30 min, 90 min and 5 h at room temperature. The samples were then briefly washed twice with de-ionised water to remove any salts and immediately plunge-frozen by immersing into liquid nitrogen-cooled ethane. Samples were then freeze-dried in a liquid nitrogen-cooled turbo freeze-drier (Quorum K775X). After that, the samples were coated with 15 nm iridium using a Quorum K575X sputter coater. The images were taken by FEI Verios 460 scanning electron microscope run at 1 - 2 keV accelerating voltage. Secondary electron images were acquired using a high-resolution Through-Lens detector in full immersion mode.

*Micropipette Aspiration*

To generate micropipettes, we heat and pull 1 mm glass capillaries (World Precision Instrument) with a micropipette puller (P97, Sutter Instrument) and cut the tapered tip with a microforge (MF830, Narishige) to an inner diameter of 4 - 10 μm. Glass pipettes are plasma treated for one minute at high RF level (PDC-32G plasma cleaner, Harrick Plasma). Then the micropipette tips are passivated with PEG-silane (Gelest) overnight at room temperature. The sample chamber for micropipette aspiration is a 360 μm-deep chamber comprised of three double-sided sticky spacers (Life Technologies) stacked together and then sandwiched by two pieces of No. 1.5 coverslips (Corning). To insert the micropipette tip to the chamber we cut a 2 mm x 360 μm opening on the side of the silicone spacer. To prevent condensates from wetting the coverslips, the surfaces of coverslips are plasma treated and then PEG-silane coated using the same methods as for the micropipette tips. To generate FUS condensates, we dilute a concentrated FUS stock solution with water. The final solution contains 1.5 μM FUS, 2.5 mM Tris, 50 mM KCl, 0.05 mM DTT, 0.25% glycerol, pH 7.4. Micropipette aspiration is conducted on a Leica SP5 confocal microscope equipped with a home-built micromanipulation system. The condensates with diameters ranging from 20 μm – 40 μm are used. During each experiment, a condensate is aspirated into a suction pipette. The negative pressure in the suction pipette is applied via a vacuum controller (Fluigent) with a resolution of 0.75 Pa and a response time of 2 seconds. Several pressure steps are used for the measurement of each condensate. Videos of the aspirated condensate is recorded as the negative pressure steps between different values. Tongue length values are extracted from the videos using ImageJ. Prior to measuring the viscosity of the FUS condensates, we pre-calibrate the system with a PEG-Dextran aqueous two-phase system using the method in (*6*). Specifically, we use an aqueous solution of 5 wt % PEG 6 kDa (Sigma) and 6.4 wt % Dextran 500 kDa (Spectrum). The solution phase separates under room temperature. We measure the true viscosity of the dextran-rich and the PEG-rich phases using a rheometer (DHR-3, TA Instruments). Taking these viscosity values, we measure an M factor of 76 for our micropipette aspiration system. The M factor is used for calculating the viscosity of FUS condensate samples. The elastic modulus of the sample is calculated using the analysis in (*7*)

*Spatial Dynamic Mapping (SDM)*

By collecting time-lapse images, it is possible to analyze the time dependence of the spatial variance, $\sigma^2$, of the difference signal, *d*, obtained by subtracting the first image from each subsequent one (*8*, *9*):

$$\sigma^2(\Delta t) = \int |d(x, y; \Delta t)|^2 dxdy$$

where $d(x, y; \Delta t) = I(x, y; \Delta t) - I(x, y; 0)$, in which $I(x, y; \Delta t)$ is the intensity of each frame at time $t = \Delta t$.

We divided each frame into ROIs of about 2x2 $\mu$m$^2$, and extracted the spatial variance, $\sigma^2(\Delta t)$, from each ROI. We averaged the results of at least 100 frame pairs for each time interval and ROI, plotted the curve and extract the characteristic time from its exponential behavior. Finally, we obtained heat maps for different condensate cross-section. Calibration performed on aqueous suspension of nanoparticles confirmed the direct proportionality of the decay time versus the diffusion coefficient. By reducing the numerical aperture of the condenser lens, $NA_C$, the white light can be made partially coherent in a thin volume whose thickness, $\theta_z$, is proportional to the light wavelength, $\lambda$, and $NA_C$ via: $\theta_z = \frac{\lambda}{NA_C^2}$ so that for white light illumination and $NA_C$~0.15, $\theta_z$~20 μ$m$ (*10*–*12*). This permitted us to extract information coming only from specific regions, namely the middle and bottom plane.

*Spatial Dynamic Mapping (SDM) calibration*

We performed a series of tests to calibrate our novel SDM approach. To do that, we used the same frame sequence collected in real space for the DDM study and we divided them into ROIs and extracted for each the characteristic decay. We then built heat maps showing for each nanoparticle size the distribution of the decay time. This is supposed to be uniform within the sample although some fluctuation due to local experimental error is present. The results are shown in Fig. S2 where it is possible to appreciate the capability of this new approach to discriminate between different tracers size locally.

*Backscattering Confocal Microscopy Speckle Analysis*

The FUS condensate is formed using the same method as used for the condensates studied with micropipette aspiration. A 24-hr old condensate is imaged using the reflection mode of a point scanning confocal microscopy (Zeiss LSM 980) to collect the backscattered signal. The reflection mode uses a T80/R20 filter in the optical path, and collects the same wavelength of light as the excitation wavelength. We use a 40x water-immersion objective with a numerical aperture of 1.1, and set the pinhole size to 1 Airy unit. We use a 405-nm wavelength laser for which the range of scattering vectors that contribute to the speckle pattern is $q$ = 36.4 um$^{-1}$ to $q$ = 41.2 um$^{-1}$, where we adjust the wavelength to account for the index of refraction of water. A series of single-plane images at the equatorial plane are collected at 53 ms per frame for 1 min. The intensity correlation is calculated as discussed in (*13*). In brief, the intensity correlation function is defined using $c_I(\vec{x},t,\tau) = \frac{<I(\vec{x},t)I(\vec{x},t+\tau)>}{<I(\vec{x},t)><I(\vec{x},t+\tau)>} - 1$ , where the brackets mean averaging over nearby pixels. We determine the size of a speckle by calculating the spatial correlation of the image, and use a Gausian window whose width is this size, which is 1.6 pixels, to average the neighboring pixels in calculating the temporal correlations. The intensity correlation is normalized by the image contrast, defined as the correlation at $\tau = 0$, giving $\tilde{c_I}(\vec{x},t,\tau) = \frac{c_I(\vec{x},t,\tau)}{c_I(\vec{x},t,0)}$ . For each pixel at each lag time, we average the intensity correlation over time to obtain the time-averaged intensity correlation.

*Simulation*

FUS MINIMAL COARSE-GRAINED SIMULATIONS

We employ a minimal model for FUS proteins recently developed (*14*). in which we integrate: (1) peptide interaction binding energies of disordered *vs.* structured Low-Aromatic-Rich Kinked Segments (LARKS) from atomistic simulations (*15*), and (2) interfacial free energies and protein domain contact probabilities of FUS (full-sequence) condensates evaluated through residue-resolution coarse-grained simulations (*16*). In our minimal simulations, we model FUS proteins as 20-bead Lennard–Jones (LJ) polymers in which the different domains of FUS (PLD (residues 1–165), RGG1 (residues 166–284), RRM (residues 285–371), RGG2 (residues 372–422), ZF (residues 423–453) and RGG3 (residues 454–526)) are coarse-grained, with one bead representing ca. 26 amino acids. That is, 6 beads for the FUS-PLD, and 14 beads for the RGG1, RRM, RGG2, ZF, and RGG3 domains. A ratio of 6/20 PLD *versus* total FUS beads recapitulates the ratio of PLD *versus* total FUS amino acids (i.e., 163/526). The solvent is explicitly modelled using single-bead LJ particles that mimic water-water and water-ion interactions.

Beads that are not directly bonded to each other, establish non-bonded interactions described via the LJ potential:

$$V(r) = 4\epsilon\left[\left(\frac{\sigma}{r}\right)^{12} - \left(\frac{\sigma}{r}\right)^{6}\right]$$

where ϵ is the depth of the LJ potential, *r* the distance between two beads, and σ the molecular diameter of each

bead. The mass of every bead was chosen to be $m^* = 1$ (in reduced units). The parameters (ϵ and σ) for the various self- and cross-interactions are shown in Tables *S1* and *S2*. For convenience, we employ reduced units, which are defined as: $T^* = k_BT/\epsilon$, $\rho^* = (N/V)\sigma^3$, $p^* = p\sigma^3/\epsilon$, and time as $t^* = t/(\sigma(m/\epsilon)^{1/2})$.

The cut-off of the LJ interactions is set to 3 times the value of σ. Bonds between consecutive beads are modelled

with a harmonic potential $V_{har} = K(r - r_0)^2$ of $K = 40\ \epsilon/\sigma^2$, and a resting position of $r_0 = 1.3\sigma$. Furthermore, we apply an angular potential of the form, $Vang = K\theta\ (\theta - \theta 0)^2$, between consecutive bonds, with an angular constant of $K\theta = 0.2$ ϵ/rad$^2$ and a resting angle of $\theta0 = 180°$ for fully disordered PLD FUS replicas, and with a constant of $K\theta = 4$ ϵ/rad$^2$ and $\theta0 = 180°$ for FUS with ordered PLD, to account for the partial rigidification of the proteins after exhibiting the disorder-to-order fibrillar transition (*17*). The model has been parameterized to recapitulate the observed behaviour in FUS proteins with both disordered-like and structured-like PLD interactions (Figs. 3 and S7 of Ref. (*14*)) and to reproduce the relative density between water and FUS condensates (*18–20*).

To dynamically mimic (*21*) the structural diversity of FUS proteins during aging, we employ a time-dependent minimal coarse-grained algorithm (*14*, *17*). Based on the above minimal coarse-gained parametrization (Tables S1 and S2), we start with a system composed of a homogeneous single phase of FUS proteins with fully disordered PLDs (Fig. 5B of the main text). Then, FUS-PLD domains can spontaneously transition from their fully disordered state to the ordered state depending on the local environment. Every 100 simulation timesteps, the algorithm evaluates whether the conditions around each fully disordered FUS protein are favorable for undergoing an 'effective' disorder-to-order cross-β-fibrillar transition (the exact conditions are described below), and thus modify the protein parameters given in Tables S1 and S2, to those corresponding to ordered PLD-FUS proteins. These conditions are evaluated on the central bead of the PLD, which is a good proxy for the average position of the LARKS found in the PLD of FUS (*14*).

Our dynamic algorithm changes the identity of two FUS chains from the fully disordered state to the ordered PLD interaction parameters once the two following conditions are met: (1) Their two central beads are at a smaller distance than 2.75 σ, and (2) both central beads are surrounded, within a cut-off distance of 2.75 σ (a sensible distance close to the maximum distance at which FUS-PLD beads can still attractively interact; the potential cut-off is 3.25 σ), by at least four other central beads and six solvent particles (i.e., characteristic crowded environment of a FUS-rich liquid phase described in Ref. (*22*) through all-atom and residue-resolution simulations). Since it has been recently reported (via PMF atomistic simulations) that strong protein binding between structured FUS LARKS occurs after undergoing a disorder-to-cross-

β-sheet transition (typically of the order of 20 to 45 $k_BT$; see Ref. (*14*)), we set the transition towards ordered-PLD FUS replicas to be irreversible. To carry out these simulations, we employ the USER-REACTION (*23*) package of LAMMPS (*24*) which allows us to change the topology of the underlying system components on-the-fly.
The criterion that at least four peptides should be in close contact to trigger a disorder-to-order transition has been chosen based on the following arguments. Four interacting peptides is the minimal system where all the different types of stacking and hydrogen bonding interactions that stabilize the β-sheet fibrillar ladder are fulfilled; i.e., two interacting steps of the ladder each made of a pair of β-sheet peptides. Thus, considering fewer interacting peptides (e.g., only three or two) would severely underestimate the required interaction strength among ordered LARKS to form irreversible inter-peptide b-sheet motifs, and subsequently, erroneously preclude the formation of kinetically arrested states at the coarse-grained level. Atomistic simulations from Refs. (*14*, *17*, *25*) show that the strength of interactions among four ordered peptides is already high enough for the peptides to remain stably bound upon thermal fluctuations. Hence, if we made the criterion even more stringent (i.e., by requiring clustering of five or more peptides), the strength of interactions among the system would remain consistent with kinetic arrest at the coarse-grained level. However, a stringent criterion would now render the coarse-grained simulations prohibitively expensive. That is, much longer simulation timescales would be needed to capture the rarer higher-density fluctuations that could result in the spontaneous formation of clusters of five or more peptides (as opposed to the more frequent fluctuations that yield clusters of four peptides; as shown in Fig. 5c of the main text).

The system size of our FUS minimal simulations (Fig. 5 of the main text) contained 12300 solvent particles and 1088 protein replicas. NVT simulations were run at $T^* = 3.5$ and at density of $p^* = 0.25$. Temperature was kept constant with a Nosé–Hoover thermostat (*26*) and with a relaxation time of $t^* = 0.4$. The Verlet equations of motion are integrated with a timestep of $t^* = 0.0004$. We employ the Direct Coexistence method as described in Refs. (*18*, *27*, *28*) and the LAMMPS Molecular Dynamics software (*24*).
**Table S1**: Model parameters for the different ϵ values in reduced units.

| | Solvent | PLD ordered | PLD disordered | Non-PLD ordered | Non-PLD disordered |
|---|---|---|---|---|---|
| Solvent | 3.50 | 2.20 | 1.07 | 1.15 | 1.15 |
| PLD ordered | 2.20 | 3.80 | 1.20 | 1.50 | 1.20 |
| PLD disordered | 1.07 | 1.20 | 1.45 | 1.25 | 1.25 |
| Non-PLD ordered | 1.15 | 1.50 | 1.25 | 1.50 | 1.25 |
| Non-PLD disordered | 1.15 | 1.20 | 1.25 | 1.25 | 1.25 |

**Table S2**: Model parameters for the different σ values in reduced units.

| | Solvent | PLD ordered | PLD disordered | Non-PLD ordered | Non-PLD disordered |
|---|---|---|---|---|---|
| Solvent | 1.00 | 1.15 | 1.15 | 1.15 | 1.15 |
| PLD ordered | 1.15 | 1.30 | 1.30 | 1.30 | 1.30 |
| PLD disordered | 1.15 | 1.30 | 1.30 | 1.30 | 1.30 |
| Non-PLD ordered | 1.15 | 1.30 | 1.30 | 1.30 | 1.30 |
| Non-PLD disordered | 1.15 | 1.30 | 1.30 | 1.30 | 1.30 |

References

1. T. Murakami, S. Qamar, J. Q. Lin, G. S. K. Schierle, E. Rees, A. Miyashita, A. R. Costa, R. B. Dodd, F. T. S. Chan, C. H. Michel, D. Kronenberg-Versteeg, Y. Li, S.-P. Yang, Y. Wakutani, W. Meadows, R. R. Ferry, L. Dong, G. G. Tartaglia, G. Favrin, W.-L. Lin, D. W. Dickson, M. Zhen, D. Ron, G. Schmitt-Ulms, P. E. Fraser, N. A. Shneider, C. Holt, M. Vendruscolo, C. F. Kaminski, P. St George-Hyslop, ALS/FTD Mutation-Induced Phase Transition of FUS Liquid Droplets and Reversible Hydrogels

into Irreversible Hydrogels Impairs RNP Granule Function. *Neuron*. **88**, 678–690 (2015).
2. S. Qamar, G. Wang, S. J. Randle, F. S. Ruggeri, J. A. Varela, J. Q. Lin, E. C. Phillips, A. Miyashita, D. Williams, F. Ströhl, W. Meadows, R. Ferry, V. J. Dardov, G. G. Tartaglia, L. A. Farrer, G. S. Kaminski Schierle, C. F. Kaminski, C. E. Holt, P. E. Fraser, G. Schmitt-Ulms, D. Klenerman, T. Knowles, M. Vendruscolo, P. St George-Hyslop, FUS Phase Separation Is Modulated by a Molecular Chaperone and Methylation of Arginine Cation-π Interactions. *Cell*. **173**, 720-734.e15 (2018).
3. Y. Shen, F. S. Ruggeri, D. Vigolo, A. Kamada, S. Qamar, A. Levin, C. Iserman, S. Alberti, P. S. George-Hyslop, T. P. J. Knowles, Biomolecular condensates undergo a generic shear-mediated liquid-to-solid transition. *Nature Nanotechnology*. **15**, 841–847 (2020).
4. F. S. Ruggeri, G. Longo, S. Faggiano, E. Lipiec, A. Pastore, G. Dietler, Infrared nanospectroscopy characterization of oligomeric and fibrillar aggregates during amyloid formation. *Nature Communications*. **6**, 1–9 (2015).
5. F. S. Ruggeri, B. Mannini, R. Schmid, M. Vendruscolo, T. P. J. Knowles, Single molecule secondary structure determination of proteins through infrared absorption nanospectroscopy. *Nature Communications*. **11**, 2945 (2020).
6. H. Wang, F. M. Kelley, D. Milovanovic, B. S. Schuster, Z. Shi, Surface tension and viscosity of protein condensates quantified by micropipette aspiration. *Biophysical Reports*. **1**, 100011 (2021).
7. D. P. Theret, M. J. Levesque, M. Sato, R. M. Nerem, L. T. Wheeler, The Application of a Homogeneous Half-Space Model in the Analysis of Endothelial Cell Micropipette Measurements. *Journal of Biomechanical Engineering*. **110**, 190–199 (1988).
8. M. Linsenmeier, M. Hondele, F. Grigolato, E. Secchi, K. Weis, P. Arosio, Dynamic arrest and aging of biomolecular condensates are modulated by low-complexity domains, RNA and biochemical activity. *Nature Communications*. **13**, 3030 (2022).
9. R. Cerbino, V. Trappe, Differential Dynamic Microscopy: Probing Wave Vector Dependent Dynamics with a Microscope. *Physical Review Letters*. **100**, 188102 (2008).
10. M. Riccomi, F. Alberini, E. Brunazzi, D. Vigolo, Ghost Particle Velocimetry as an alternative to μPIV for micro/milli-fluidic devices. *Chemical Engineering Research and Design*. **133**, 183–194 (2018).
11. T. Pirbodaghi, D. Vigolo, S. Akbari, A. DeMello, Investigating the fluid dynamics of rapid processes within microfluidic devices using bright-field microscopy. *Lab on a Chip*. **15**, 2140–2144 (2015).
12. S. Buzzaccaro, E. Secchi, R. Piazza, Ghost Particle Velocimetry: Accurate 3D Flow Visualization Using Standard Lab Equipment. *Physical Review Letters*. **111**, 048101 (2013).
13. S. Aime, M. Sabato, L. Xiao, D. A. Weitz, Dynamic Speckle Holography. *Physical Review Letters*. **127**, 088003 (2021).
14. A. Garaizar, J. R. Espinosa, J. A. Joseph, G. Krainer, Y. Shen, T. P. J. Knowles, R. Collepardo-Guevara, Aging can transform single-component protein condensates into multiphase architectures. *Proceedings of the National Academy of Sciences*. **119** (2022), doi:10.1073/pnas.2119800119.
15. P. Robustelli, S. Piana, D. E. Shaw, Developing a molecular dynamics force field for both folded and disordered protein states. *Proc Natl Acad Sci U S A*. **115**, E4758–E4766 (2018).
16. J. A. Joseph, A. Reinhardt, A. Aguirre, P. Y. Chew, K. O. Russell, J. R. Espinosa, A. Garaizar, R. Collepardo-Guevara, Physics-driven coarse-grained model for

biomolecular phase separation with near-quantitative accuracy. *Nature Computational Science 2021 1:11*. **1**, 732–743 (2021).

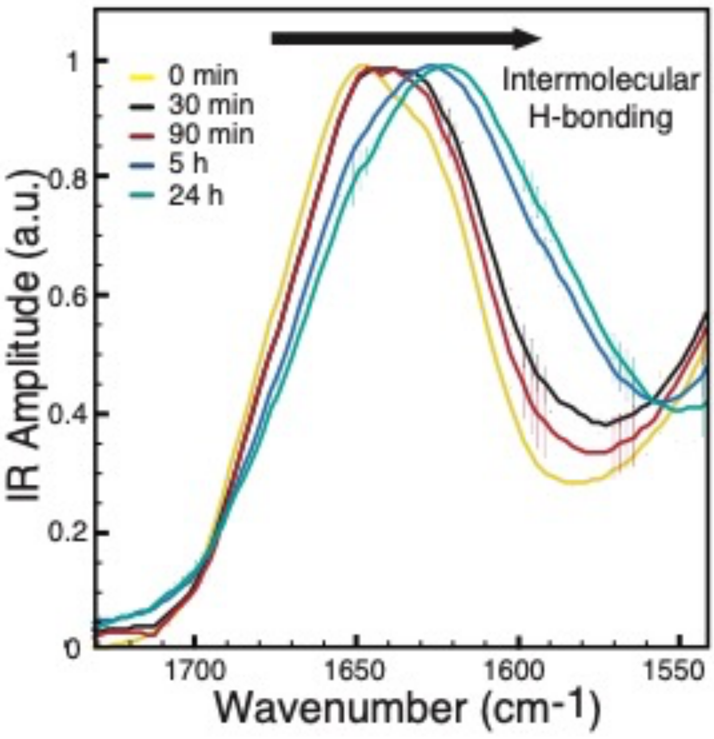


Fig. S1, The shift of the wavenumber in IR spectra of the FUS condensates incubated for different time at room temperature.

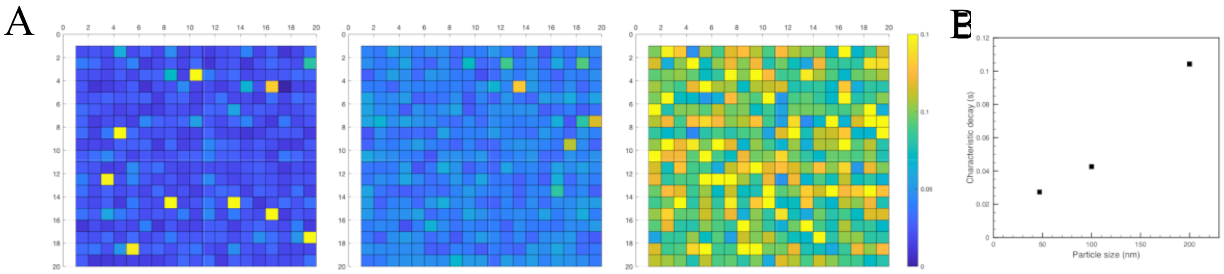


Fig. S2. Nanoparticle calibration of the system. A Spatial Dynamic Mapping (SDM) for 47, 100 and 200 nm particle respectively showing the uniform distribution of the characteristic decay time (values in seconds). B Average characteristic decay time obtained from SDM versus particle size. The error bar (standard error) is smaller than the symbol.
